# nf-core/detaxizer: A Benchmarking Study for Decontamination from Human Sequences

**DOI:** 10.1101/2025.03.27.645632

**Authors:** Jannik Seidel, Camill Kaipf, Daniel Straub, Sven Nahnsen

**Author notes:** Joint last authors (Daniel Straub, Sven Nahnsen).

## Abstract

Privacy is paramount in health data, particularly in human genetics, where information extends beyond individuals to their relatives. Metagenomic datasets contain substantial human genetic material, necessitating careful handling to mitigate data leakage risks when sharing or publishing. The same applies to genetic datasets from the environment or datasets from contaminated laboratory samples, although to a lesser extent. To address these topics, we developed nf-core/detaxizer, a nextflow-based pipeline that employs Kraken2 and bbmap/bbduk for taxonomic classification, identifying and optionally filtering *Homo sapiens* reads. Due to its generalized design, other taxa can also be classified and filtered. We benchmark its filtering efficacy for human reads against Hostile and CLEAN, demonstrating its utility for secure data preprocessing. The comparison revealed that the choice of tool and database can lead to an order of magnitude more human data not removed. As part of the nf-core initiative, nf-core/detaxizer adheres to best practices, leveraging containerized dependencies for streamlined installation. The source code is openly available under the MIT license: https://github.com/nf-core/detaxizer.

## INTRODUCTION

Human sequencing data can be found in various samples, from environmental sources as bycatch (1) to directly human-associated data. In Whole-Genome Sequencing (WGS) metagenomic data originating from a human being the amount of human reads varies depending on the site of origin. For example, up to 10% human reads can be found in stool samples, while in saliva samples they can exceed 90% (2). This raises ethical and data privacy related concerns as a person can be identified with a certain probability from such data (3). Thus, for efficiently researching microbial communities based on metagenomics data, all traces from human data must be removed. To date, even the most sophisticated available filtering techniques miss to remove these traces comprehensively. Absolute human-free data cannot be achieved due to interindividual variability and the fact that one simply cannot construct a database that covers the complete genetic variability within mankind. Also, sequences shared between *Homo sapiens* and other taxa cannot be distinguished. To advance current decontamination capabilities from metagenomic data, nf-core/detaxizer was developed in nextflow (4) within the nf-core framework (5) to allow reproducibility and easy usage. A modular approach was pursued with nf-core/detaxizer to integrate several tools because a combination of different approaches of human read removal performed best (6). nf-core/detaxizer is presented in this manuscript and compared to two other tools for filtering, namely CLEAN (7) and Hostile (8). In this study, we show that all tested tools performed well, but the most thorough removal of human sequences was achieved by nf-core/detaxizer. Decontamination performance and alleviating data privacy concerns varied by magnitudes, exacerbated by database choices.

Due to the generalized design of nf-core/detaxizer, other taxa can also be classified and filtered, therefore this benchmark can also guide separating all kinds of taxa from sequencing data.

## MATERIAL AND METHODS

### nf-core/detaxizer

The pipeline nf-core/detaxizer was implemented using nextflow (4), a Domain Specific Language (DSL) in the field of bioinformatics and data science for orchestrating workflows, in version 2 within the nf-core framework (5).

Optionally validation via blastn can be performed. The output of the labelling and optional validation is summarized. Filtering is performed optionally and the decontaminated taxonomic profile can be inspected with Kraken2.

The workflow in version 1.1.0 (Fig. 1) utilizes Kraken2 2.1.3 (9) and/or bbmap/bbduk 39.10 (https://sourceforge.net/projects/bbmap/) (hereafter termed as bbduk) for classification of the reads in fastq format. Both tools allow the customization of the databases used in the classification step.

**Figure 1.**
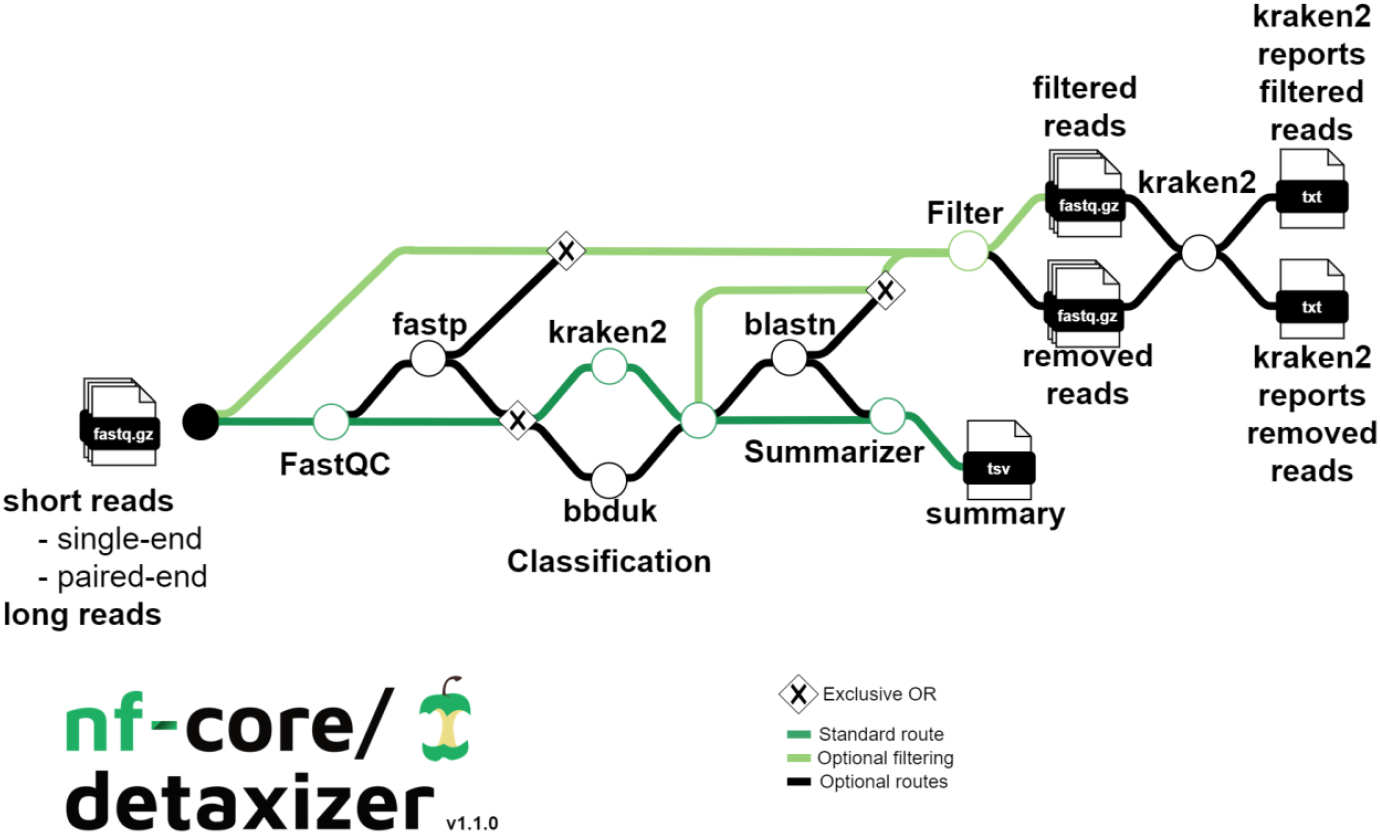
Overview of the nf-core/detaxizer workflow (from left to right). Input can be short and/or long reads. Quality control is performed via FastQC and optionally the reads can be pre-processed via fastp. By default, Kraken2 labels reads for removal and optionally bbmap/bbduk can be chosen. The output of labelled reads is merged if both tools are used.

For filtering data with Kraken2, a model was developed which allows the fine-tuning of the classification step. First, a taxonomic name (e.g. “*Homo sapiens*”) via the ‘tax2filter’ parameter is provided that is supposed to be identified. For a non-leaf-node of the taxonomic tree the whole subtree is labelled. The classification logic has three conditions that all need to be fulfilled to label a read (pair): (1) The number of k-mers of the designated taxonomy must be above a threshold (parameter ‘cutoff_tax2filter’), (2) the ratio of the number of k-mers of the designated taxonomy to the number of all other k-mers that are not ‘unclassified’ plus the number of k-mers of the designated taxonomy must be above a threshold (parameter ‘cutoff_tax2keep’), and (3) the ratio of the number of k-mers of the designated taxonomy to the number of ‘unclassified’ k-mers plus the number of k-mers of the designated taxonomy must be above a cutoff (parameter ‘cutoff_unclassified’). To use bbduk, a fasta file with the sequences of the respective taxon or taxa to label must be provided via the ‘fasta_bbduk’ parameter. The k-mer size used in the classification process can be set, having a default value of 27 (parameter ‘bbduk_kmers’). bbduk produces fastq files of reads with at least one k-mer assigned to the provided fasta. These fastq files are then transformed into a list of labelled read (pair) IDs using seqkit 2.8.2 (10).

The union of labelled read (pair) IDs by Kraken2 and bbduk is considered final, if both classifiers are used, and blastn 2.15.0 (11) can be further used to remove potential false positives. An overview with the number of labelled read (pairs) per tool and sample serves as a summary of filtering efforts. If preprocessing with fastp (12) is performed by setting parameter ‘preprocessing’, pre-processed files instead of raw read files can be filtered by setting the ‘enable_filter’ parameter. This optional step employs seqkit 2.8.2 (10) to remove all labelled sequences from the respective fastq files. The removed sequences can be optionally copied into the results folder via the ‘output_removed_reads’ parameter.

### Benchmark

#### Dataset

The dataset (13) used for the benchmark had 10,000,002 human read pairs and 21,172,961 microbial read pairs. The human reads (NovaSeq 6000, paired-end 150bp) originated from 3 human individuals (Finish male; Luhya, Kenya male; Bengali, Bangladesh female) from the 1000 Genomes Project. The microbial reads contained data derived from ZymoBIOMICS HMW DNA Standard D6322 (Zymo Research, seven bacterial strains and one fungal; MiSeq, paired-end 151bp; termed Zymo from this point onward) and *Mycobacterium tuberculosis* (HiSeq 4000, paired-end 75bp; termed MTB from this point onward).

#### nf-core/detaxizer settings

Performance of nf-core/detaxizer with combinations of Kraken2, bbduk, and various databases was benchmarked. Details are shown in Table 1. The parameters of the Kraken2 k-mer model (‘cutoff_tax2filter’, ‘cutoff_tax2keep’, ‘cutoff_unclassified’) were set to 0 to label as many human reads as possible. These settings correspond to labelling a read pair as human contamination if at least one k-mer was assigned to human irrespective of k-mer matches to other taxa. Increasing these settings will reduce the number of read pairs labelled as human contamination. To perform the benchmarking, nextflow was installed on a workstation in version 24.04.4 and Apptainer (14) as container engine in version 1.3.5-1.fc40 to utilize BioContainers (15). The exact command used for the benchmark can be found in Supplementary 1. Parameter combinations are shown in Table 1.

**Table 1:**
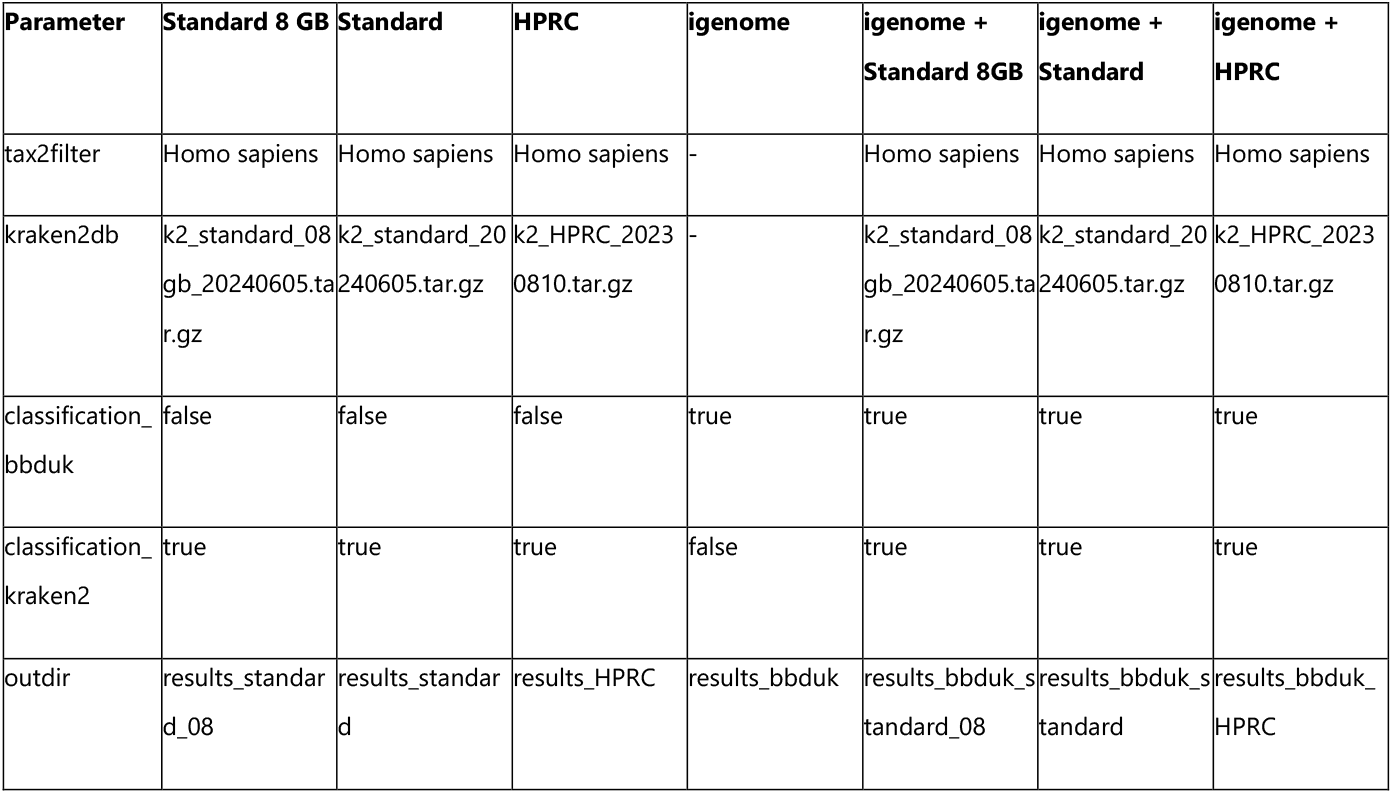
Parameter combinations for nf-core/detaxizer benchmarking. All analyses also included parameters ‘enable_filter’, ‘output_removed_reads’, and ‘input samplesheet.csv’. Parameters set but without effect on labelling were ‘classification_kraken2_post_filtering’ and ‘save_intermediates’. bbduk automatically uses GRCh38 AWS igenome (termed igenome); other fasta files can be provided via the ‘fasta_bbduk’ parameter. Details about the databases can be found in Supplementary 1.

#### Hostile

Hostile 1.1.0 (8) was installed via conda 24.7.1. It used bowtie2 2.4.5 (16) for classification. The default database was used, consisting of the t2t human genome plus HLA sequences (8). The exact commands used for the benchmark can be found in Supplementary 1.

#### CLEAN

CLEAN 1.0.3 (7) is based on nextflow and employs for short read decontamination bbduk 38.79. The databases used were ‘hsa’ and ‘t2t’ provided by CLEAN via the ‘autodownload’ function. Also, the GRCh38 AWS igenome was used to investigate whether the performance of CLEAN (using bbduk 38.79) and nf-core/detaxizer with bbduk 39.10 was identical. The exact command used for the benchmark can be found in Supplementary 1.

#### Evaluation of decontaminated Data

Human data (read pairs flagged by the authors of the benchmarking dataset as “human”) was deemed contamination. False positives (FP) and true negatives (TN) were microbial, true positives (TP) and false negatives (FN) human read pairs. FP and TP were removed, TN and FN appeared in decontaminated data. The read ID extraction and counting was performed with Python 3.12.4 (17) and modules: gzip, re, Biopython 1.78 (18) with SeqIO.

Recall and precision (19) were calculated to gain an overview of how well the tools decontaminated the data. Recall was the measure for the success of the decontamination (1).

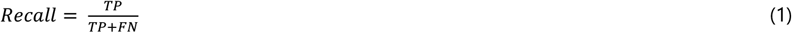

Precision indicated how well microbial reads were retained (2).

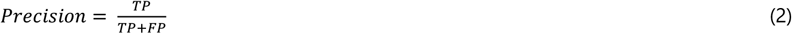

To determine if the ratio of the two categories, Zymo and MTB, in the two resulting datasets (TN in the decontaminated reads and FP in removed reads) were dependent upon the categories in the original dataset or not a 2×2 χ^2^-test of independence (20) was performed for each outcome.

## RESULTS

The benchmarking of combinations of pipelines and databases for human decontamination of sequencing data revealed generally high performance but also striking differences. Missed human read pairs varied by a factor of up to 18.5 (from 3,770 to 69,725 FN) and wrongly removed sequences by 152 times (from 728 to 110,992 FP).

In terms of recall, the best performing decontamination was achieved by nf-core/detaxizer with the combination of bbduk with GRCh38 AWS igenome and Kraken2 with the Kraken2 Standard database (Fig. 2A). nf-core/detaxizer reached a recall of 0.99962 and missed only 3,770 human read pairs (FN). Of these 3,770 FN, 98.17% (3,701) were unclassified by Kraken2 with the Kraken2 Standard database. The remaining 69 FN (1.83%) were annotated by Kraken2 as *archaea* (1), *viruses* (2), or *bacteria* (65), and 1 FN as cellular organism. Among those 65 FN annotated as *bacteria*, 58 were attributed with a phylum, of which 28 were classified as *Pseudomonadota* (including 7 *Ralstonia*) and 16 as *Bacillota* (including 5 *Clostridium*). Notably, the labelling by nf-core/detaxizer with Kraken2 is done on the k-mer level, while the taxonomic classification by Kraken2 is a consensus of all k-mer matches. Of 85,741 erroneously removed read pairs (FP), 69,686 (81.32%) originated from the Zymo dataset and 16,055 (18.68%) from the MTB dataset. The benchmarking dataset contained similar read numbers for Zymo and MTB, 46.92% and 53.08%, respectively. Therefore, wrongly decontaminated microbial reads were strongly skewed by 34.4 percentage points towards the Zymo dataset (2×2 χ^2^-test, p < 0.001). The retained microbial reads (TN) with 9,864,089 (46.78%, input: 46.92%) Zymo and 11,223,131 (53.22%, input: 53.08%) MTB read pairs were also significantly skewed by 0.14 percentage points towards MTB (2×2 χ^2^-test, p < 0.001).

**Figure 2.**
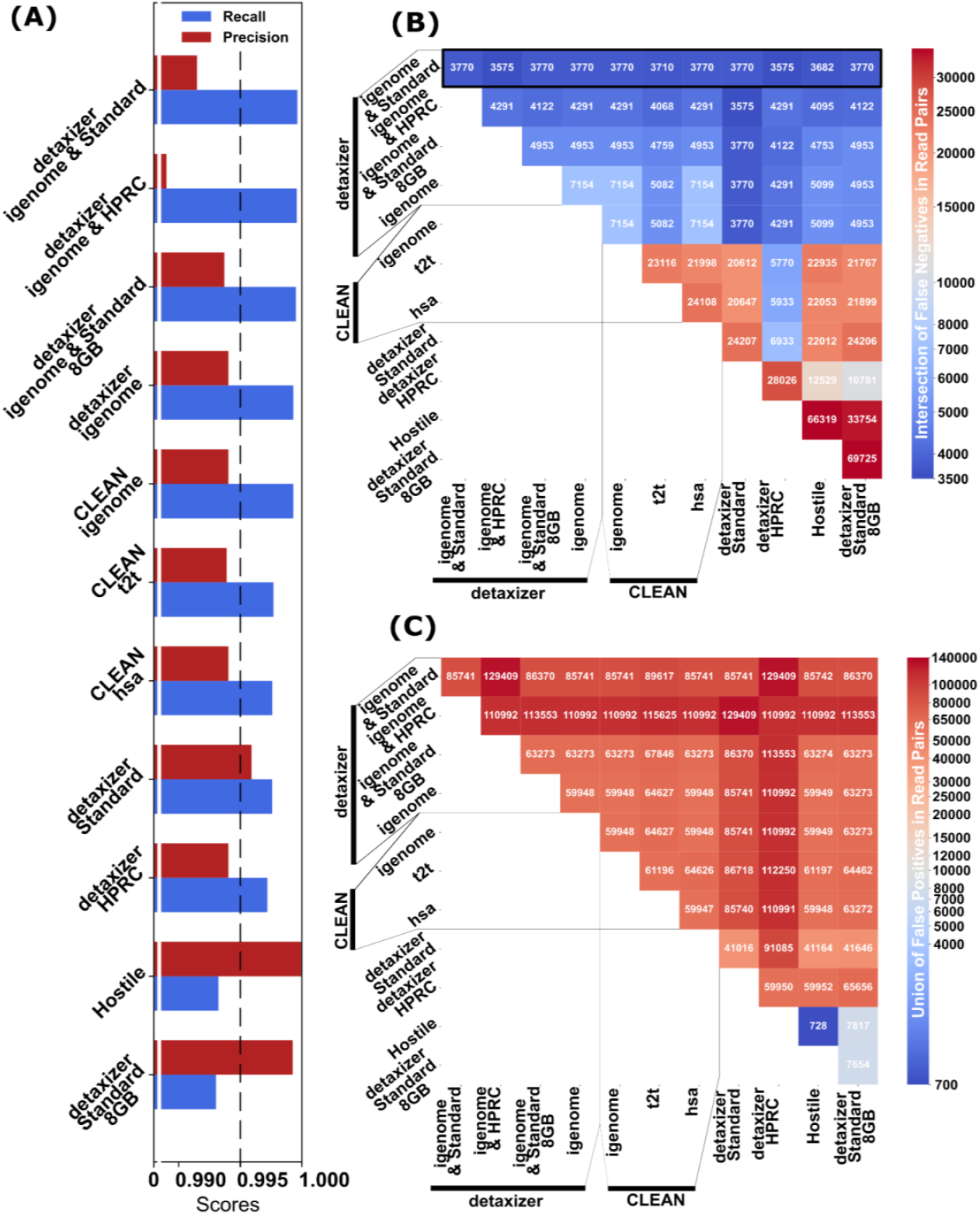
Decontamination performance of pipeline and database combinations. Pipelines were nf-core/detaxizer, CLEAN, and Hostile. Tools for classification were Kraken2, bbduk and bowtie2. Databases were Kraken2 Standard, HPRC, Kraken2 Standard 8GB, GRCh38 AWS igenome, t2t, hsa, default of Hostile. (A) Recall and precision of pipelines and selected database(s). Data is sorted from top to bottom by decreasing recall, i.e. top is the most thorough decontamination solution. (B) Number of not detected human read pairs (false negatives) for intersections between pipelines and settings. Data is sorted decreasing from top to bottom. The best performing decontamination of a single pipeline missed 3,770 read pairs (nf-core/detaxizer with bbduk and Kraken2 with Standard database). The most thorough decontamination missed 3,575 read pairs and is achieved by chaining two methods in 3 combinations. (C) Union of microbial read pairs labelled wrongly as human contamination (false positives). The input was approx. 10 million human and approx. 21 million microbial read pairs.

Whether a combination of tools would increase recall was examined by intersecting the FN read pairs (Figure 2B) and uniting the FP read pairs (Figure 2C) of all examined tools. The combination of “nf-core/detaxizer bbduk with GRCh38 AWS igenome and Kraken2 with Kraken2 Standard database” with “nf-core/detaxizer HPRC” reduced FN by 125 read pairs and increased recall slightly from 0.99962 to 0.99964 compared to the best performing combination in Figure 2A but increased the amount of FP substantially by 51% (additional 43,668 read pairs) thereby reducing precision from 0.99150 to 0.98722.

Hostile performed best regarding precision (0.99992, equals 728 FP) with a recall of 0.99322 (66,319 FN) (Figure 2A). nf-core/detaxizer with Kraken2 Standard 8GB database followed closely with precision of 0.99922 (7,654 FP) and recall of 0.99303 (69,725 FN), i.e. 0.0007 and 0,00019 less than Hostile for precision and recall, respectively.

## DISCUSSION

To discuss the importance of filtering human reads from metagenomic data, one must acknowledge that a re-identification can be performed with a certain probability when the complete genome of the data donor is available (3). The re-identification probability depends on the site of origin of the sample, the number of reads, and the coverage compared to the human genome. Given the number of reads is the same, the likelihood of re-identification can be estimated to be higher for a sample from saliva with more than 90% of human reads (2) than from stool with up to 10% human reads. The dependence upon the whole genetic information of the data donor reduces the real-world risk to a certain extent, though it cannot be discarded. Human reads could be extracted using a method like nf-core/detaxizer and be used for further analysis, revealing characteristics of the donor like sex, ethnicity (3) and potentially genetic diseases. Combining multiple data points facilitates identification and might be used to harm the data donor, and by extension potentially relatives.

Decontamination from human reads in (meta)genomics is a non-trivial task which is of utmost importance to preserve data privacy. Based on the presented benchmark, nf-core/detaxizer v1.1.0 is recommended with the Kraken2 Standard database together with bbduk with the GRCh38 AWS igenome. Blastn was considered to reduce false positives but was rejected in this study because of high time requirements and increase of false negatives which is undesirable when aiming for data privacy. Hostile has 10 times lower false positives and is recommended in cases when a bias of the remaining microbial reads has to be valued higher than data privacy. Though, a skew of 0.14% towards MTB in the TN data with nf-core/detaxizer is likely irrelevant for subsequent microbial abundance analyses. The bias might be due to the different sequencing length of MTB and Zymo, PE75 and PE151, respectively. This statement should be further validated on other high-quality datasets with a known ground truth and higher taxonomic diversity, but a general recommendation can be derived that filtering of human reads should not influence downstream microbial analyses while preserving data privacy to a high extent. Also, it was shown with simulated metagenomic data with human contamination that analyses are significantly different when comparing pre- and post-filtering datasets (21). Taken together, further studies with more benchmarking datasets could benefit the research on how different filtering methods perform and how decontamination improves analyses.

Another question arises when looking at the false negative results, as most are unclassified. One could argue that all unclassified reads should be filtered from the data to maximize data privacy. This would lead to a near perfect recall. But an additional 225,559 (compared to 85,741) unclassified microbial reads would be false positives in this case. Filtering must be balanced between recall and precision and the presented setting for nf-core/detaxizer seems to maximize recall while not abandoning precision. It can be speculated that human reads not identified in this study as human might be either genuine contamination of the human sequencing data, sequencing artefacts or highly divergent sequences from e.g. variable region (HLA) or proviruses.

To conclude, filtering of human reads from metagenomic data is inevitable when publishing or sharing such data to protect the data donor’s privacy to the extent possible, however, no available method seems perfect, and a slight bias in the decontaminated microbial data cannot be excluded at that point.

## ACKNOWLEDGEMENTS

We would like to thank the nf-core team for welcoming us to the nf-core community and their assistance in reviewing and releasing the pipeline. We would also like to thank the nf-core community for developing the nf-core infrastructure and resources for Nextflow pipelines. A full list of nf-core community members is available at https://nf-co.re/community.

ChatGPT 4 was utilized in developing nf-core/detaxizer and 4o to enhance the writing style of this manuscript.

## AUTHOR CONTRIBUTIONS

Jannik Seidel: Conceptualization, Formal analysis, Methodology, Software, Validation, Visualization, Writing—original draft. Camill Kaipf: Methodology. Daniel Straub: Conceptualisation, Supervision, Writing—original draft. Sven Nahnsen: Supervision, Writing—review & editing.

## SUPPLEMENTARY DATA

Supplementary Data are available online.

## DATA AVAILABILITY

nf-core/detaxizer code is hosted on GitHub under the nf-core organization https://github.com/nf-core/detaxizer and at Zenodo https://doi.org/10.5281/zenodo.10877147 and is released under the MIT license. The version 1.1.0 is hosted at Zenodo (https://doi.org/10.5281/zenodo.14056601).

